# Phosphatidylinositol 5-phosphate 4-kinase (PIP4K) regulates sugar homeostasis in *Drosophila*

**DOI:** 10.64898/2025.12.03.692006

**Authors:** Arnab Karmakar, Padinjat Raghu

## Abstract

Metabolic adaptation such as sugar homeostasis is central to the response of living organisms to environmental changes and in insects, sugar homeostasis is an integral part of larval metabolism. Mutants of phosphatidylinositol 5-phosphate 4-kinase (dPIP4K) show altered growth and larval development. We find that *dPIP4K* mutants (*dPIP4K^29^)* show enhanced glucose uptake by the fat body, accompanied by increased circulating trehalose levels in feeding third instar larvae. Following acute starvation on a low sugar diet, *dPIP4K^29^* larvae show reduced survival associated with high circulating trehalose levels, while under equivalent conditions, wild-type larvae undergo depletion of circulating trehalose. While Tret1-1PA transcripts were upregulated on starvation in muscle and brain tissue of controls, this enhancement was abrogated in *dPIP4K^29^*. Muscle specific depletion of Tret1-1PA in wild type larvae prevents reduction of circulating trehalose levels upon starvation, phenocopying that seen in *dPIP4K^29^*and selective reconstitution of dPIP4K in muscle of *dPIP4K^29^* could rescue starvation responses to wild type levels. Together, these findings identify dPIP4K as a key regulator of sugar homeostasis in larval tissues and survival in larvae during starvation.

## Introduction

Phosphoinositides are signalling lipids present at very low abundance in cellular membranes. They play important roles as second messengers regulating multiple aspects of eukaryotic cell function including G-protein coupled receptor and receptor tyrosine kinase signalling, vesicular transport, cytoskeletal organization, ion channel and transporter activity (Balla, 2013). Based on the positions at which the inositol ring is phosphorylated, there are seven phosphoinositide species in cells. The generation and turnover of these lipids is mediated by enzymes, the phosphoinositide kinases and phosphatases (Sasaki et al., 2009). These enzymes are highly selective with respect to the substrate they utilise as well as the position of the hydroxyl group on the inositol headgroup that they phosphorylate. Therefore, these enzymes are central to the ability of phosphoinositides to regulate cell and animal physiology. Phosphatidylinositol 5-phosphate 4-kinase (PIP4K) is one such kinase, which converts phosphatidylinositol 5 phosphate (PI5P) into phosphatidylinositol 4,5 bisphosphate [PI(4,5)P_2_] [(Rameh et al., 1997)]. Like other phosphoinositide kinases, PIP4K also exhibits exquisite substrate specificity; it is able to convert PI5P into PI(4,5)P_2_ but does not utilise phosphatidylinositol 4 phosphate (PI4P) as a substrate [reviewed in (Krishnan et al., 2026)]. Interestingly, genes encoding PIP4K are typically found only in genomes of multi-cellular organisms and are not found in unicellular eukaryotes (Krishnan et al., 2025). While genomes of invertebrates such as *Drosophila* and *C. elegans* contain a single PIP4K gene, in mammalian genomes, three isoforms exist, namely, PIP4K2A, PIP4K2B, and PIP4K2C. Investigation using mouse models has revealed the importance of these enzymes in animal physiology; loss of each isoform in mouse results in specific phenotypes. For example, loss of both PIP4K2A and PIP4K2B slows down tumour growth in p53^-/-^ mice (Emerling et al., 2013) while depletion of PIP4K2C results in excessive T cell activation (Shim et al., 2016).

A number of studies have suggested a key role for PIP4K in regulating metabolism. In rodent models, depletion of PIP4K2B results in hyper-responsiveness to insulin and a progressive loss of body weight in adults (Lamia et al., 2004). Likewise, in *Drosophila*, loss of the only PIP4K (*dPIP4K*) results in a larval growth deficit and developmental delay, which is associated with an overall reduction of target of rapamycin complex 1 (TORC1) activity (Gupta et al., 2013). Interestingly, *dPIP4K* mutant larvae show enhanced Class I phosphatidylinositol 3 kinase (Class I PI3K) activity, as indicated by increased phosphatidylinositol 3,4,5 trisphosphate (PIP_3_) levels at the plasma membrane of salivary glands and fat body cells (Sharma et al., 2019). This enhanced activation of Class I PI3K in response to receptor tyrosine kinase stimulation was also recapitulated in human cells when PIP4K genes were deleted (Wang et al., 2019). Moreover, loss of *dPIP4K* protects against the metabolic consequences of a high sugar diet (HSD). When reared on a HSD, wild type larvae develop hypertrehalosemia and show delayed development (Musselman et al., 2011). By contrast, *dPIP4K* mutant larvae, upon rearing on a HSD, show unchanged levels of circulating trehalose and a less severe developmental delay compared to control flies under the same conditions (Sharma et al., 2019). These two observations-reduced growth and increased insulin sensitivity in *dPIP4K^29^*are paradoxical, since enhanced insulin signalling in larval tissues is expected to increase larval growth rate. One possible explanation is that *dPIP4K* has distinct functions in various larval tissues or specific impacts on the regulation of particular aspects of metabolism. In *Drosophila,* growth and development is controlled through the activity of multiple larval tissues (Droujinine and Perrimon, 2016) and signalling pathways (Ahmad et al., 2020). Key among these are the fat body that regulates both fat and sugar metabolism (Meschi and Delanoue, 2021) and insulin producing cells of the brain that produce insulin like peptides (Nässel and Broeck, 2016).

The fat body regulates lipid metabolism but is also a key tissue in the regulation of sugar metabolism; it plays a key role in the synthesis and release of trehalose which is the main circulating sugar in *Drosophila* haemolymph. The fat body is the *Drosophila* counterpart of the liver and adipose tissue of mammals and plays pivotal roles in regulating sugar and lipid metabolism in flies. A key function of the larval fat body is to maintain sugar homeostasis, regulated by insulin signalling (Arrese and Soulages, 2010). Fat body cells uptake glucose from the circulation and rapidly converts it into trehalose and glycogen (Arrese and Soulages, 2010; Li et al., 2019). Trehalose, being the main circulating sugar, is released into the circulation by the fat body via trehalose transporter Tret1-1 (Kikawada et al., 2007), whereas glycogen is stored inside fat cells. Trehalose metabolism plays a pivotal role in enabling larval adaptation to diverse nutritional conditions (Yasugi et al., 2017). Under starvation, circulating trehalose is taken up by peripheral tissues and metabolised as an energy source; this is critical for larval survival (Tellis et al., 2023) and the function of individual organs during starvation, a process referred to as organ sparing (e.g brain sparing).

In this study, we analysed sugar metabolism in *Drosophila* larvae and found that *Drosophila dPIP4K* mutant larvae have altered sugar homeostasis. This was manifested as enhanced glucose uptake by the fat body, associated with increased circulating trehalose levels compared to controls. Further, *dPIP4K^29^* larvae maintain high levels of circulating trehalose even during nutrient starvation in contrast to wild type animals that show starvation induced reduction in the circulating trehalose levels. We found that the likely reason for this phenotype is the loss of starvation response in *dPIP4K^29^*. Under acute starvation, circulating trehalose utilisation is enhanced in selected tissues including the glial cells of the brain and muscle, mediated by upregulation of trehalose transporter Tret1-1PA, a process that is lost in *dPIP4K^29^*. Consequently, larval survival and adaptation under starvation is impaired in *dPIP4K^29^*. Overall, we find dPIP4K to be important for maintaining sugar homeostasis, loss of which affects larval response to nutrient deprivation.

## Results

### Loss of *dPIP4K* alters sugar homeostasis in larvae

Since it was previously reported that the fat body of *dPIP4K^29^*showed enhanced insulin sensitivity (Sharma et al., 2019), we wondered how this would impact larval metabolism and physiology. In mammals, insulin signalling activates GLUT4 mediated glucose uptake in muscle and adipose tissue; an equivalent process is reported to exist in the *Drosophila* fat body, although the ortholog of GLUT4 in flies has not been identified (Crivat et al., 2013). Since insulin sensitivity is increased in the fat body of *dPIP4K^29^*(Sharma et al., 2019), we checked if glucose uptake is altered in feeding third instar larvae. The rate of glucose uptake was tested by dissecting and incubating fat body *ex vivo* for 15 min with 2-[N-(7-nitrobenz-2-oxa-1,3-diazol-4-yl)amino]-2-deoxy-D-glucose (2-NBDG), a fluorescent glucose analogue, followed by imaging to estimate 2-NBDG uptake (Hirabayashi et al., 2013). We found that the level of 2-NBDG uptake was significantly increased in the fat body of *dPIP4K^29^* larvae compared to wild type **(Fig 1A,B)**. However, it has been reported that 2-NBDG can be metabolized after uptake inside cells, while 6-(*N*-(7-Nitrobenz-2-oxa-1,3-diazol-4-yl)amino)-6-Deoxyglucose (6-NBDG), another fluorescent-glucose analogue, is non-metabolizable (Yamamoto et al., 2011). Hence, to confirm if the increased signal of 2-NBDG in *dPIP4K^29^*is due to increased uptake and not by reduced metabolism inside cells, the level of uptake was tested using both 2-NBDG and 6-NBDG; we found that uptake of both compounds was enhanced in *dPIP4K^29^* compared to wild type **(Fig S1A,B)**. These findings suggest that glucose uptake is enhanced in fat body cells of *dPIP4K^29^*.

**Figure 1.**
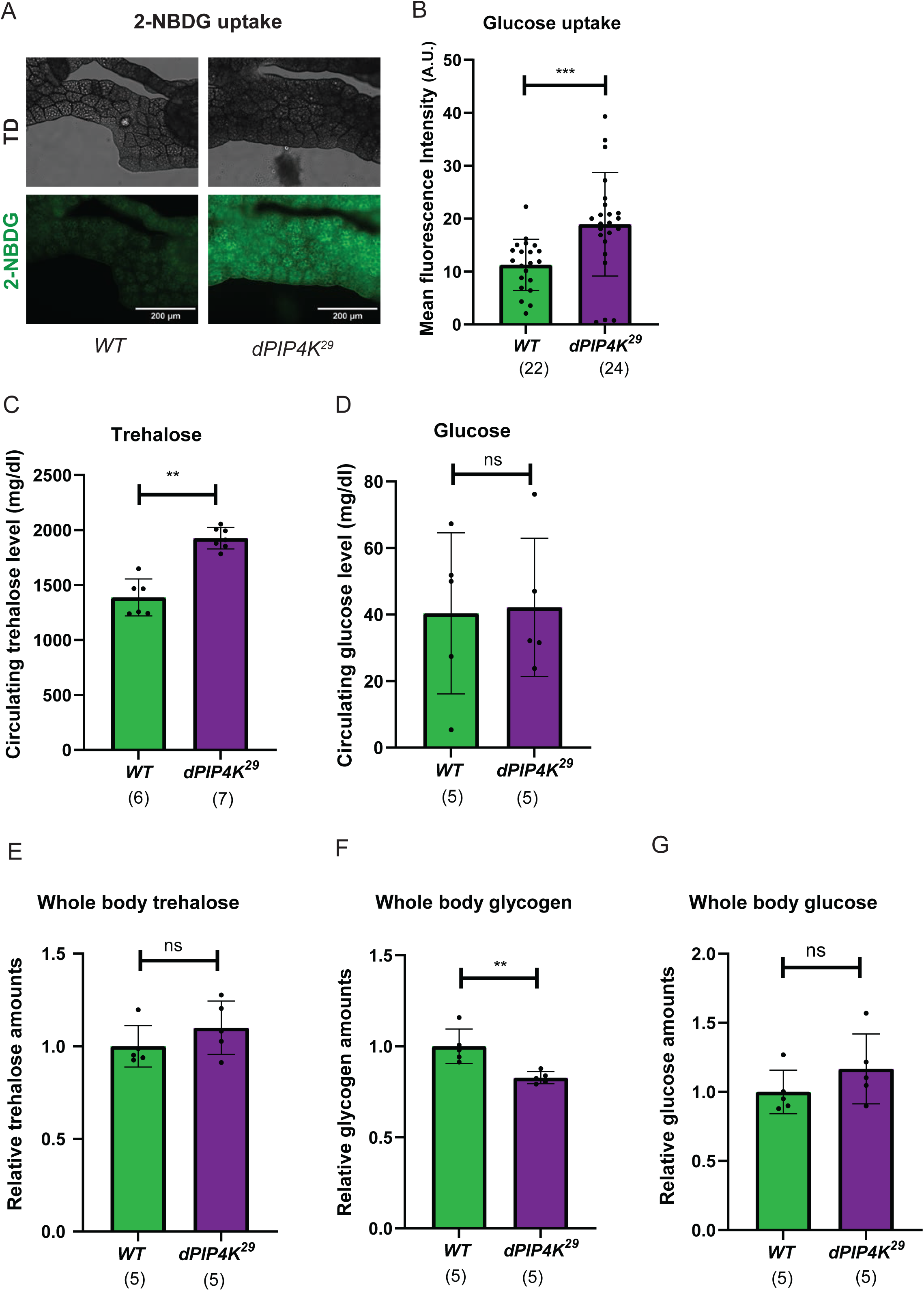
Altered sugar homeostasis in *dPIP4K^29^* larvae. **(A)** Representative images and **(B)** quantification of 2-NBDG uptake in the fat body of wild type (*WT*) and *dPIP4K^29^* feeding third instar larvae. Brightfield images (TD) are shown in the upper panels and the corresponding fluorescent images depicting 2-NBDG uptake on lower panels are shown (Scale bar: 200 μm). Each data point in the graph is one biological replicate representing the mean fluorescence intensity (in arbitrary unit, A.U.) of the fat body collected from a single larva. Measurements of **(C)** circulating trehalose level and **(D)** circulating glucose level from the haemolymph of *WT* and *dPIP4K^29^* larvae. Each data point is one biological replicate representing average circulating trehalose measured from pooled haemolymph of 5-8 feeding third instar larvae. Relative amounts of **(E)** trehalose, **(F)** glycogen, and **(G)** glucose from the whole larval extracts of *WT* and *dPIP4K^29^*. Each data point represents measured values of the respective metabolites normalized to protein, and further normalized to the corresponding mean values of *WT*. For (**B-G**), each bar represents the mean ± SD. The numbers of biological replicates are indicated in parentheses below each bar. Mann-Whitney U test was used for the statistical analysis*. **p-value < 0.01, ***p-value < 0.001, ns-*non-significant *p-value > 0.05*.

We analysed the levels of glucose and trehalose from haemolymph and from whole larval extracts of feeding 3^rd^ instar larvae. The level of circulating trehalose, the major circulating sugar in flies, was elevated in *dPIP4K^29^* **(Fig 1C)** while that of glucose was unchanged compared to control **(Fig 1D)**. On the other hand, the levels of trehalose **(Fig 1E)** and glucose **(Fig 1G)** from whole larval extracts were unchanged, while the level of glycogen was marginally reduced in *dPIP4K^29^*compared to controls **(Fig 1F)**. These findings indicate altered sugar metabolism in *dPIP4K^29^*.

The fat body is also the major lipid storage organ in fly larvae (Arrese and Soulages, 2010). Lipid is stored in the form of triglycerides (TAG) inside lipid droplets (LDs) in the fat body, which actively go through lipogenesis or lipolysis depending on nutrient availability. Changes in lipid metabolism can be detected by analysing the size of LDs in the fat body (Musselman et al., 2011). Using Nile red staining, we compared the size of fat body LDs between *dPIP4K^29^* and wildtype, and found that there was no difference **(Fig S1C)**. In addition, a biochemical assay for total triglycerides showed no difference in levels between wild type and *dPIP4K^29^*either in whole body extracts or the fat body **(Fig S1D,E)**. Overall, these finding suggest that there is a defect in sugar homeostasis but not lipid metabolism in *dPIP4K^29^*.

### *dPIP4K^29^* is resistant to depletion of circulating trehalose levels during starvation

Since trehalose is utilised as an energy source during starvation, we checked whether the starvation response in terms of circulating trehalose is altered in *dPIP4K^29^*. Trehalose metabolism acts as a metabolic buffer for larval development during nutrient shortage (Matsushita and Nishimura, 2020). During starvation, circulating trehalose is utilised by peripheral tissues as an energy source, where it is broken down into glucose by the action of cytoplasmic trehalase (cTreh) (Matsuda et al., 2015; Tellis et al., 2023). Since feeding larvae of *dPIP4K^29^* have enhanced circulating trehalose levels, we tested how nutrient starvation affects their circulating trehalose pool. We found that in wild type feeding third instar larvae, starvation led to the rapid depletion of circulating trehalose levels within 6 hrs, consistent with a recent report (González-Gutiérrez et al., 2025), and this reduction of trehalose levels was enhanced with longer periods of starvation up to 24 hrs; however, under the same conditions, there was a much smaller reduction of trehalose levels in *dPIP4K^29^* larvae **(Fig 2A,B)**. Pan-larval depletion of dPIP4K led to a smaller depletion of circulating trehalose levels upon starvation than controls (**Fig 2C, S2A)** partially phenocopying *dPIP4K^29^*. Reconstitution of dPIP4K in *dPIP4K^29^*could partially rescue starvation induced circulating trehalose depletion to that seen in wild type **(Fig 2D, S2B)**. Together, these findings suggest PIP4K regulates circulating trehalose levels in *Drosophila* larvae during starvation.

**Figure 2.**
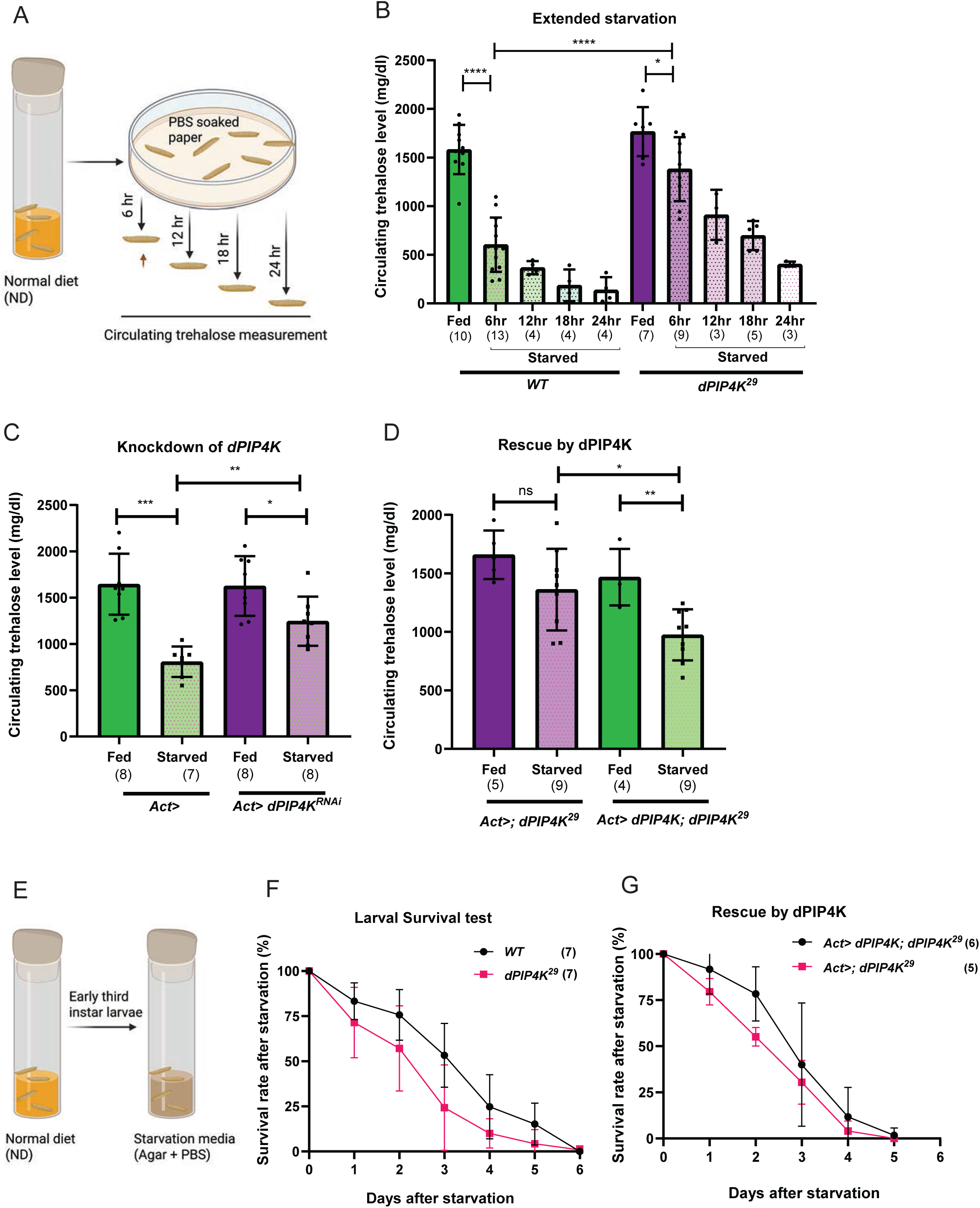
Starvation response is impaired in *dPIP4K^29^* larvae. **(A)** Schematic overview of the experimental design for circulating trehalose measurements following larval starvation of four time points-6, 12, 18, and 24 hrs in PBS soaked filter paper. **(B)** Quantification of circulating trehalose level from the feeding and starved larvae (four time points) of *WT* and *dPIP4K^29^*. For all further trehalose measurement under complete starvation experiments, 6 hrs starvation timepoint was considered (shown by arrowhead in **Fig 2A**). Comparison of circulating trehalose levels between fed and starved larvae of the following genotypes-**(C)** *Act>* (control) and *Act> dPIP4K^RNAi^* (depletion of dPIP4K), and **(D)** *Act>; dPIP4K^29^* (mutant control) and *Act> dPIP4K; dPIP4K^29^*(dPIP4K reconstitution). (**E)** Schematic representation for larval survival test after starvation, for which early third instar larvae were transferred to starvation media (0.8% agar in PBS), and the number of larvae alive post starvation were counted every 24 hrs afterwards. The larval survival rate is shown for the genotypes-**(F)** *WT* and *dPIP4K^29^*, and **(G)** *Act>; dPIP4K^29^*(mutant control) and *Act> dPIP4K; dPIP4K^29^* (dPIP4K reconstitution). Y-axis represents the percentage of larvae alive and X-axis shows time in days post larval transfer to starvation media (Day 0 is the day of transfer). For (**B,C,D**), each data point is one biological replicate, representing average circulating trehalose measured from pooled haemolymph of 5-8 larvae. Each bar represents the mean ± SD. The numbers of biological replicates are indicated in parentheses below each bar. Mann-Whitney U test was used for the statistical analysis*, *p-value < 0.05, **p-value < 0.01, ***p-value < 0.001, ****p-value < 0.0001, ns-*non-significant *p-value > 0.05*.

Alterations in trehalose homeostasis by loss of trehalose 6 phosphate synthase (Tps1) or trehalase (Treh) impairs larval adaptation to various dietary conditions or nutrient deprivation (Yasugi et al., 2017). Hence, we tested whether the altered trehalose utilisation by *dPIP4K^29^*affects their larval adaptation and survival. Indeed, the survival of early third instar larvae was reduced in *dPIP4K^29^* compared to wild type upon starvation **(Fig 2E,F)**, and partially rescued by reconstitution of dPIP4K **(Fig 2G)**.

We also examined if sugar-specific starvation is sufficient to cause comparable alterations of circulating trehalose and larval survival. For this, a low-sugar diet (LSD) was prepared with yeast extract and agar, without any carbohydrate sources **(Fig S2C)**. Early-third instar larvae were transferred onto LSD for 24 hrs, which caused a significant reduction in the circulating trehalose level of wild type, but not of *dPIP4K^29^* **(Fig S2D)**. Further, larval survival rate after transferring them onto LSD was reduced in *dPIP4K^29^* compared to wildtype **(Fig S2E)**, recapitulating the impact of total starvation **(Fig 2F)**. By contrast, the impact of amino-acid starvation on circulating trehalose is no different between wildtype and *dPIP4K^29^*, as larvae of both genotypes exhibited no change in circulating trehalose levels upon feeding on a no-protein diet (NPD) for 24 hrs **(Fig S2F,G)**. Overall, loss of dPIP4K prevents trehalose depletion in the circulation under nutrient restrictions, affecting larval adaptation and response to starvation.

### Trehalose synthesis and release during starvation is not altered in *dPIP4K^29^*

How is starvation induced reduction of circulating trehalose lost in *dPIP4K^29^* larvae? It has been reported that during starvation, peripheral tissues such as glia perform increased uptake of trehalose from the circulating haemolymph, which is utilised as an energy source to ‘spare’ the brain from starvation (Hertenstein et al., 2021; Volkenhoff et al., 2015) leading to depletion of the circulating trehalose pool (González-Gutiérrez et al., 2025). On the other hand, circulating trehalose can be replenished by the fat body via increased breakdown of glycogen into glucose and its conversion into trehalose which is then release into the haemolymph **(Fig 3A)** (Yamada et al., 2018). In principle, the failure of starvation induced depletion of circulating trehalose in *dPIP4K^29^* could result from either increased trehalose production and release into haemolymph by the fat body, or reduced trehalose uptake from the circulation during starvation.

**Figure 3.**
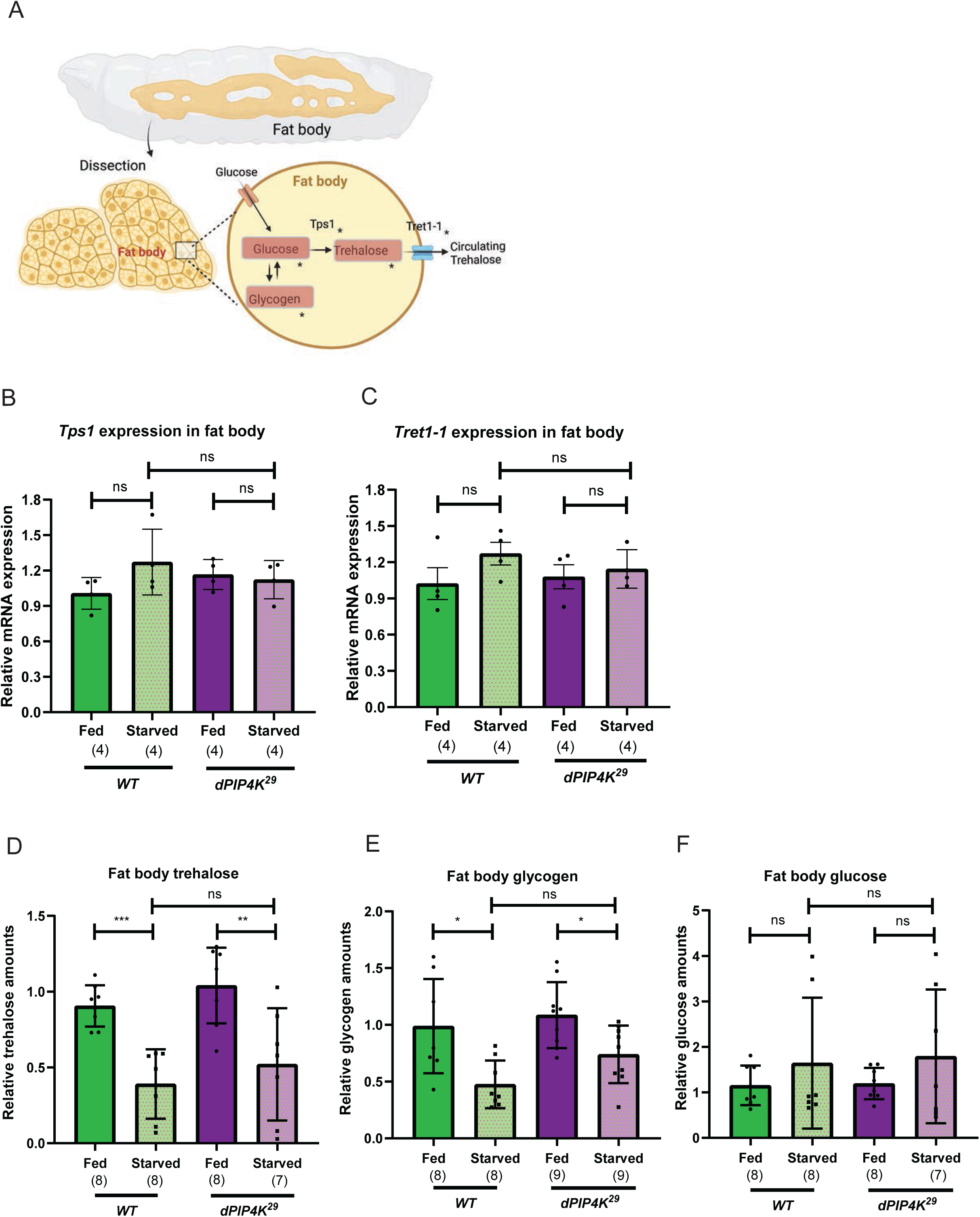
Starvation induced replenishment of trehalose in circulation is not altered in dPIP4K^29^. **(A)** Schematic representation of trehalose synthesis by fat body in a larva. The fat body from the larvae were dissected for measurement of the fat body-specific metabolites and enzymes, marked with asterisk (*). The relative mRNA expression levels of **(B)** *Tps1* and **(C)** *Tret1-1,* measured by qPCR from the fat body extracts of fed and starved *WT* and *dPIP4K^29^* larvae. Relative amounts of **(D)** trehalose, **(E)** glycogen and **(F)** glucose in fat body from fed and starved *WT* and *dPIP4K^29^* larvae. Each data point represents measured values of the respective metabolites normalized to protein, and further normalized to the corresponding mean values of *WT* fed group. For **(B-F)**, each bar represents the mean ± SD. The numbers of biological replicates are indicated in parentheses below each bar. Mann-Whitney U test was used for the statistical analysis*, *p-value < 0.05, **p-value < 0.01, ***p-value < 0.001, ns-*non-significant.

To test if trehalose synthesis and release from the fat body is increased in *dPIP4K^29^*, we compared the mRNA expression of the fat body specific trehalose synthesizing enzymes, Tps1 (Yoshida et al., 2016), and trehalose transporter, Tret1-1 that is proposed to release trehalose from the fat body into haemolymph (Kanamori et al., 2010; Tellis et al., 2023). We found that while the levels of *Tps1* transcripts were modestly elevated in wild type larval fat body on starvation, no elevation was seen in *dPIP4K^29^***(Fig 3B)**. A similar observation was noted for *Tret1-1* transcripts in fat body, which were not elevated on starvation in *dPIP4K^29^***(Fig 3C)**. Further, biochemical analysis revealed that the levels of trehalose in the fat body following starvation were no different in *dPIP4K^29^* compared to wild type **(Fig 3D)**. The levels of glycogen, that is broken down into glucose to produce trehalose under starvation were no different between wild type and *dPIP4K^29^***(Fig3E)** following starvation and the levels of fat body glucose were also not different between the two strains **(Fig 3F)**. Since there might also be other non-fat body sources of trehalose synthesis, we also estimated the levels of total trehalose, glycogen and glucose from the whole larval extracts of *dPIP4K^29^* and wild type; these were no different between the two genotypes upon starvation **(Fig S3D,E,F)**. Likewise, whole body expressions of *Tps1*, *Tret1-1* and *cTreh* were unchanged between wild type and *dPIP4K^29^* larvae under starvation **(Fig S3A,B,C).** Further, the activity of secreted Trehalase (sTreh) that breaks down circulating trehalose into glucose in the haemolymph was analysed and was no different in *dPIP4K^29^* than wild type in either fed or starved conditions **(Fig S3G)**.

Overall, these results suggest that the synthesis and replenishment of circulating trehalose upon starvation is not elevated in *dPIP4K^29^* larvae; hence is unlikely to cause the trehalose accumulation phenotype in *dPIP4K^29^*.

### Starvation induced upregulation of Tret1-1PA is lost in *dPIP4K^29^*

Next, we tested the hypothesis that the failure of trehalose uptake by tissues **(Fig 4A)** might underlie the unchanged circulating trehalose levels of *dPIP4K^29^* during starvation. Under starvation, uninterrupted supply of energy is ensured for tissues such as the brain by several mechanisms (Cheng et al., 2011); one such mechanism is via increased uptake of sugar by glial cells from circulation by increased activity of the trehalose transporter Tret1-1PA (Hertenstein et al., 2021). The mRNA expression of *Tret1-1PA* is reported to be upregulated specifically in glia, but not in neurons (Hertenstein et al., 2021). Hence, starvation induced depletion of circulating trehalose levels in larvae could be due to increased uptake of trehalose by glial cells or other tissues with high energy demand. We depleted Tret1-1 in neurons and glia independently; while neuron-specific depletion of Tret1-1 in larvae caused a drop in circulating trehalose levels upon starvation comparable to wild type flies **(Fig S4A)**, glia-specific depletion of Tret1-1 blocked trehalose depletion **(Fig 4B)**, phenocopying that seen in *dPIP4K^29^*. Under similar conditions, depletion of Tret1-1 in the fat body was without effect **(Fig S4B)**. We also performed Tret1-1 depletion in muscle, since flight muscles in adult flies extensively utilise circulating trehalose for energy production (Becker et al., 1996). Tret1-1 depletion in muscle also resulted in no depletion of circulating trehalose under starvation **(Fig 4C)**, phenocopying *dPIP4K^29^*.

**Figure 4.**
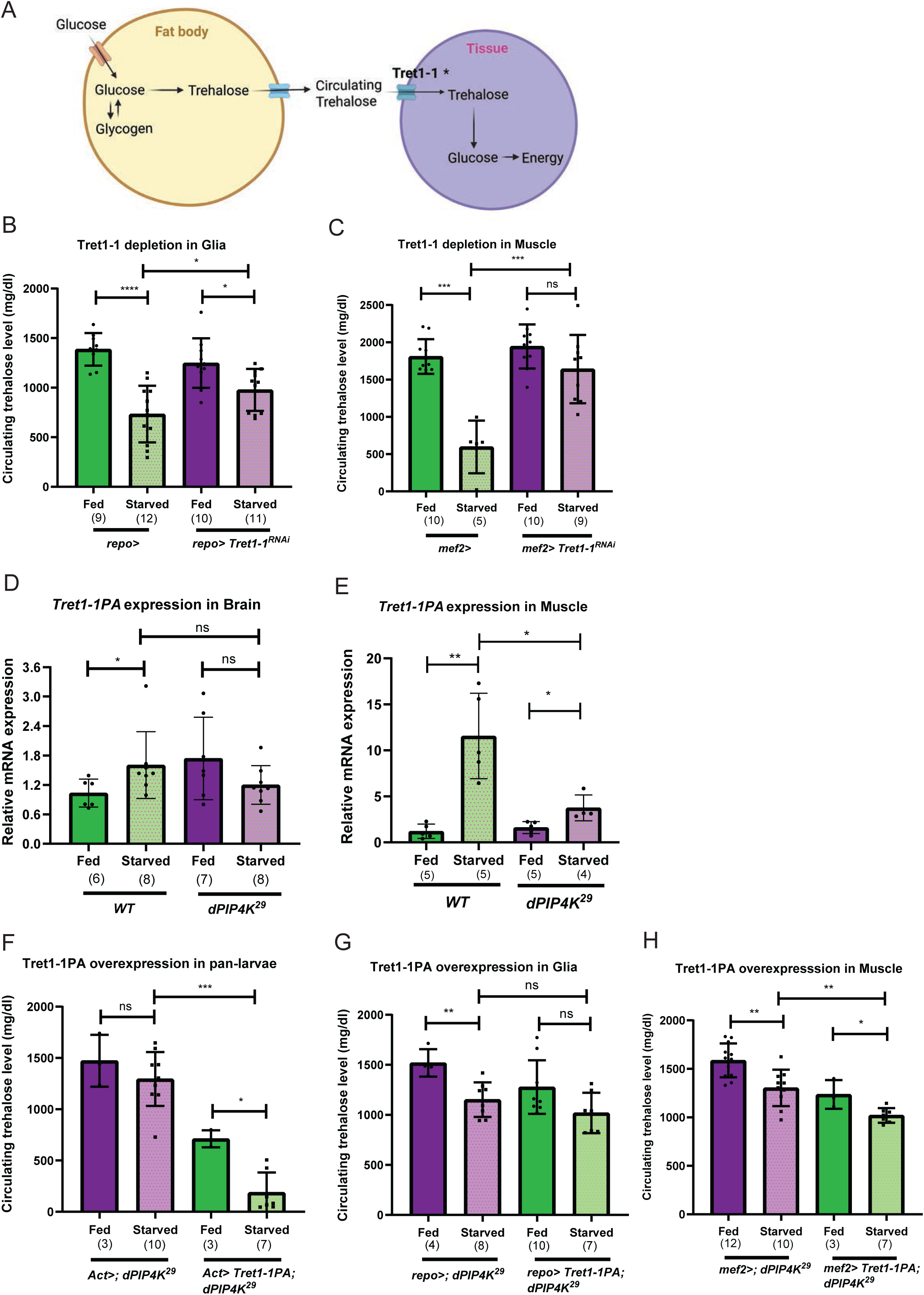
Starvation induced upregulation of *Tret1-1* is impaired in *dPIP4K^29^*. **(A)** Schematic representation depicting molecular and biochemical elements of circulating haemolymph and trehalose uptake by trehalose transporter Tret1-1 (marked with asterisk*). Measurement of circulating trehalose level following tissue-specific depletion of Tret1-1 in- **(B)** glia and **(C)** muscle. The comparison of circulating trehalose levels are shown between fed and starved conditions of the corresponding genotypes - **(B)** *repo>* (control) and *repo> Tret1-1^RNAi^* (depletion in glia) and **(C)** *mef2>* (control) and *mef2> Tret1-1^RNAi^* (depletion in muscle). The relative mRNA expression levels of *Tret1-1PA* measured by qPCR from the following tissues-**(D)** brain and **(E)** muscle of *WT* and *dPIP4K^29^* larvae under fed and starved conditions. Measurement of circulating trehalose levels following overexpression of Tret1-1PA in-**(F)** pan-larvae, **(G)** glia and **(H)** muscle of *dPIP4K^29^*. The comparison of circulating trehalose levels are shown between fed and starved conditions of the corresponding genotypes-**(F)** *Act>; dPIP4K^29^* (mutant control) and *Act> Tret1-1PA; dPIP4K^29^* (Tret1-1PA overexpression in whole larva), **(G)** *repo>; dPIP4K^29^* (mutant control) and *repo> Tret1-1PA; dPIP4K^29^* (Tret1-1PA overexpression in glia) and **(H)** *mef2>; dPIP4K^29^* (mutant control) and *mef2> Tret1-1PA; dPIP4K^29^* (Tret1-1PA overexpression in muscle). For (**B,C,F,G,H**), each point is one biological replicate, which represents the average circulating trehalose level measured of the haemolymph pooled from 5-8 larvae. For all graphs, each bar represents the mean ± SD. The numbers of biological replicates are indicated in parentheses below each bar. Mann-Whitney U test was used for the statistical analysis*, *p-value < 0.05, **p-value < 0.01, ***p-value < 0.001, ****p-value < 0.0001, ns-*non-significant *p-value > 0.05*.

Since glia and muscle specific Tret1-1 depletion phenocopied *dPIP4K^29^*, the levels of this gene product in these tissues could be altered in *dPIP4K^29^*, preventing circulating trehalose utilisation under starvation. To test this, the mRNA expression of *Tret1-1PA* was compared between brain samples of fed and starved larvae of wild type and *dPIP4K^29^*. Upon starvation, the brain-specific expression of *Tret1-1PA* was significantly enhanced in wild type, consistent with a previous report (Hertenstein et al., 2021); however the expression of this transcript in brain was unchanged in *dPIP4K^29^***(Fig 4D)**. Interestingly, the brain-specific mRNA expression of *Tret1-1PA* in the fed condition itself was significantly higher in *dPIP4K^29^* than controls and did not increase any further under starvation **(Fig 4D)**. Likewise, although *Tret1-1PA* expression was strongly upregulated in wild type muscle tissue on starvation, only a very modest upregulation was seen in *dPIP4K^29^***(Fig 4E)**. This suggests that the upregulation of *Tret1-1PA* under starvation is impaired in *dPIP4K^29^* and hence, depletion of circulating trehalose is much lower than in controls. As a test of this hypothesis, overexpression of Tret1-1PA in *dPIP4K^29^* should result in trehalose depletion upon starvation as seen in wild type animals. Indeed, pan-larval overexpression of Tret1-1PA **(Fig S4C)** led to a reduction of circulating trehalose levels in *dPIP4K^29^* under starvation **(Fig 4F)**. Under our experimental conditions, selective overexpression of Tret1-1PA in glia of *dPIP4K^29^*only led to a modest if any, reduction of circulating trehalose levels **(Fig 4G)** but a more substantial reduction was seen when Tret1-1PA was selectively overexpressed in muscle **(Fig 4H)**.

### dPIP4K function in selected tissues underlies altered sugar homeostasis

To test whether dPIP4K has a role in selected tissues for regulating trehalose homeostasis, we carried out a tissue-screen where dPIP4K was depleted individually in specific tissues involved in sugar metabolism and changes in circulating trehalose level in response to starvation were analysed **(Fig S5)**. While no single tissue-specific depletion fully recapitulated the phenotype of *dPIP4K^29^*, the starvation-induced decline in trehalose level seen in controls was partially attenuated upon glia, muscle, and neuron-specific depletion of dPIP4K compared to controls **(Fig 5A,C and S5C)**, suggesting that dPIP4K activity in multiple tissues likely contributes to the systemic regulation of circulating trehalose level. Given our observations on the effects of Tret1-1PA depletion in muscle and glia on the control of circulating trehalose levels **(Fig 4)**, we tested the effect of selective reconstitution of dPIP4K in these two tissues. Selective reconstitution of dPIP4K in muscle **(Fig S5H)** led to a partial reversal of the stable starvation induced circulating trehalose levels in *dPIP4K^29^***(Fig 5D)**. Further, muscle-specific reconstitution of dPIP4K also rescued larval survival upon starvation in *dPIP4K^29^*larvae **(Fig 5E)**. On the other hand, selective reconstitution in glia **(Fig 5B)** led to a less effective reversal of the *dPIP4K^29^* phenotype. Overall, our findings suggest that altered *Tret1-1* expression in the muscle tissue of larvae contributes to altered trehalose homeostasis and starvation response in *dPIP4K^29^*.

**Figure 5.**
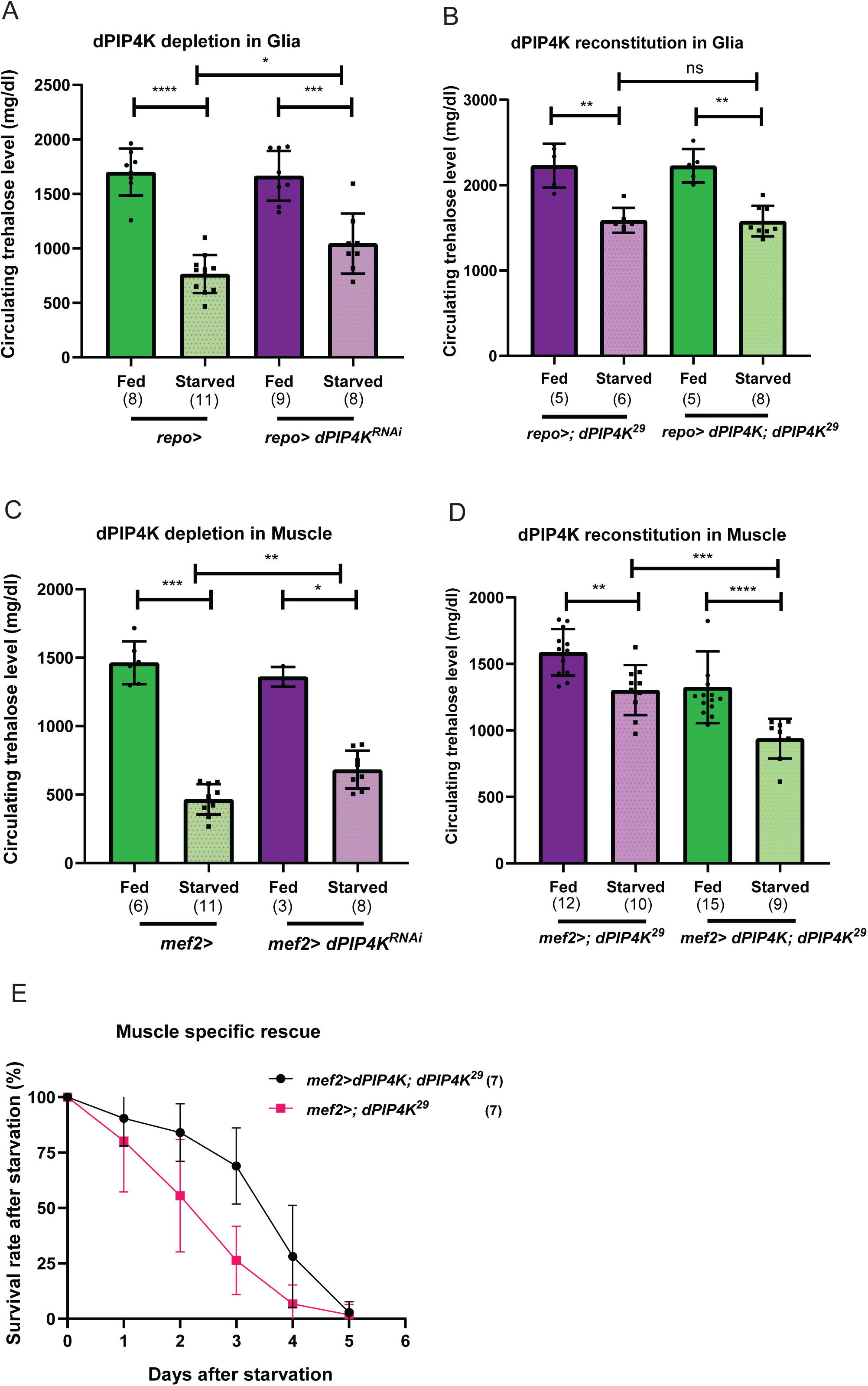
Reconstitution of dPIP4K in muscle rescues trehalose accumulation under starvation in *dPIP4K^29^*. Measurements of circulating trehalose levels upon-**(A)** glia-specific depletion of dPIP4K, **(B)** glia-specific reconstitution of dPIP4K (in mutant), **(C)** muscle-specific depletion of dPIP4K, and **(D)** muscle-specific reconstitution of dPIP4K (in mutant). The comparison of circulating trehalose levels are shown between fed and starved larvae of the corresponding genotypes-**(A)** *repo>* (control) and *repo> dPIP4K^RNAi^* (dPIP4K depletion in glia), **(B)** *repo>; dPIP4K^29^* (mutant control) and *repo>dPIP4K; dPIP4K^29^*(dPIP4K reconstitution in glia), **(C)** *mef2>* (control) and *mef2> dPIP4K^RNAi^* (dPIP4K depletion in muscle), and **(D)** *mef2>; dPIP4K^29^* (mutant control) and *mef2> dPIP4K; dPIP4K^29^*(dPIP4K reconstitution in muscle). **(E)** Larval survival rate after starvation, upon reconstitution of dPIP4K in muscle in *dPIP4K^29^* mutants. The genotypes are-*mef2>; dPIP4K^29^* (mutant control) and *mef2> dPIP4K; dPIP4K^29^* (dPIP4K reconstitution). Y-axis shows percentage of larvae alive and X-axis shows time in days post larval transfer to starvation media (Day 0 is the day of transfer). For (**A-D**), each point is one biological replicate, which represents the average circulating trehalose level measured of the haemolymph pooled from 5-8 larvae. Each bar represents the mean ± SD. The numbers of biological replicates are indicated in parentheses below each bar. Mann-Whitney U test was used for the statistical analysis*, *p-value < 0.05, **p-value < 0.01, ***p-value < 0.001, ****p-value < 0.0001, ns-*non-significant *p-value > 0.05*.

## Discussion

In multi-cellular organisms, co-ordinated regulation of nutrient homeostasis and metabolism is key to survival in response to environmental changes. For example, during changes in ambient conditions, many hormones such as insulin activate cell surface receptors which then regulate nutrient uptake into cells or the release of nutrients from cellular stores. Several such growth factors use receptors of the tyrosine kinase family that are a feature of metazoan genomes. Several receptor tyrosine kinases that activate Class I PI3K signalling are therefore key regulators of metabolism. PIP4K genes that are also restricted to metazoan genomes have emerged as regulators of receptor tyrosine kinase dependent Class I PI3K signalling (Krishnan et al., 2025). Studies in multiple metazoan models including mammals, *Drosophila* and *C. elegans* have shown that depletion of PIP4K results in metabolic defects [reviewed in (Krishnan et al., 2026)]. Insulin signalling can regulate multiple aspects of metabolism including sugars, amino acids and lipid turnover. A previous study has noted that PIP4K2B knockout mice show defects in glucose homeostasis and adiposity (Lamia et al., 2004) and reduced intestinal fat stores have been reported for the *C. elegans ppk-2* mutant (Lundquist et al., 2018). In this study we noted that *Drosophila* PIP4K mutants have defective sugar homeostasis and at a physiological level, flies depleted of PIP4K were abnormally sensitive to sugar deprivation although fat metabolism and amino acid signalling seems unaffected. Therefore, this study defines PIP4K as a key regulator of sugar homeostasis in *Drosophila* during both fed and starved conditions.

In *Drosophila*, previous work has shown that trehalose homeostasis in the circulating haemolymph depends both on its synthesis in the fat body by Tps1 and subsequent release into the haemolymph, as well as the uptake of this sugar by peripheral tissues through trehalose transporter Tret1-1 and breakdown into glucose by trehalase. Our analysis of *dPIP4K^29^*mutants indicates that enhanced trehalose synthesis is unlikely to underlie the elevated levels of circulating trehalose under starvation. Our finding that depletion of Tret1-1 in muscle and glia phenocopies *dPIP4K^29^* and that Tret1-1 mRNA levels are not upregulated by starvation in these tissues strongly suggests that the underlying defect in *dPIP4K^29^* is an inability to regulate trehalose uptake by peripheral tissues. In addition to previous studies demonstrating an important role for Tret1-1PA in glia, we find that depleting Tret1-1 in muscle phenocopied *dPIP4K^29^*. This finding may indicate that during starvation multiple larval tissues including glia, muscles and perhaps additional as yet unidentified tissues may take up trehalose to meet energy demands during starvation.

The mechanisms by which dPIP4K regulates sugar homeostasis in *Drosophila* larval cells remains to be firmly established. In this study, we found evidence of enhanced insulin stimulated glucose uptake in the fat body when dPIP4K was depleted; such cells have enhanced insulin stimulated PIP_3_ production (Sharma et al., 2019). This finding is conceptually similar to the observation that following a glucose challenge, PIP4K2B knockout mice show faster clearance of blood glucose compared to control (Lamia et al., 2004), indicating enhanced glucose uptake mechanisms. Previous studies have determined that in mammalian cells, the glucose transporter GLUT4 is translocated to the plasma membrane following insulin stimulation (van Gerwen et al., 2023). The identity of the glucose transporter that mediates glucose uptake in *Drosophila* cells remains to be discovered and identifying this transporter will enable a molecular analysis of the mechanism underlying the enhanced glucose uptake by the fat body in dPIP4K depleted cells. Despite the enhanced glucose uptake by the fat body, we could not find evidence of enhanced trehalose synthesis in the fat body and the levels of triglycerides were also not elevated. This finding is consistent with the observation (Lamia et al., 2004) in PIP4K2B knockout mice that mice on a high nutrient diet did not show increased adiposity. The mechanism underlying this remains unknown.

The synthesis of the disaccharide trehalose from glucose in the fat body is a feature of insect metabolism. Trehalose is released from the fat body and transported in the haemolymph prior to being taken up by tissues where it is then broken down and used in cellular energy metabolism. We made three key observations (i) Elevated trehalose levels in *dPIP4K* mutants could be phenocopied by down regulating Tret1-1 in muscle or glial cells (ii) Tret1-1 transcripts in muscle and brain failed to be upregulated during starvation (iii) Overexpression of Tret1-1PA in muscle could restore the starvation induced fall in haemolymph trehalose levels in *dPIP4K* mutants to that seen in controls. Together these observations strongly suggest that upon dPIP4K depletion, there is a defect in starvation induced elevation in Tret1-1 levels to facilitate uptake of trehalose into selected tissues to meet energy demand.

In insects sugar metabolism is regulated through the function of multiple tissues including the fat body, brain, midgut, endocrine organs and muscle. Insulin receptors, Class I PI3K and dPIP4K are expressed ubiquitously in fly larvae including all these tissues. Although acute, ubiquitous depletion of dPIP4K in fly larvae was able to recapitulate the abnormal trehalose homeostasis noted in germline *dPIP4K^29^* mutants, depleting dPIP4K individually in any one of these tissues was unable to phenocopy the abnormal trehalose homeostasis. This finding likely underscores the likely role of dPIP4K in multiple larval tissues in regulating sugar homeostasis.

The mechanism by which dPIP4K might regulate Tret1-1 mediated uptake of trehalose remains to be established. In *Drosophila*, dPIP4K has been implicated as a direct regulator of Insulin/Class I PI3K signalling. However, a previous study has suggested that Tret1-1 upregulation in glial cells is not dependant on insulin signalling; rather it is regulated transcriptionally by TGF-β signalling (Hertenstein et al., 2021). Therefore, it is unlikely that the defect in starvation induced upregulation of Tret1-1 is a direct consequence of the ability of dPIP4K to regulate Insulin/Class I PI3K signalling. However it is to be noted that TGF-β receptor signalling can regulate Class I PI3K activity (Hamidi et al., 2017; Yi et al., 2005) and the contribution of such regulation to Tret1-1 expression on cells remains to be tested in the context of this model system. However, Hertenstein et.al have also reported that the control of Tret1-1 expression also depends on endo-lysosomal trafficking. Interestingly, a previous study has implicated dPIP4K in the regulation of membrane protein trafficking via the endosomal system in *Drosophila* (Kamalesh et al., 2017). Therefore, it is possible that dPIP4K regulates Tret1-1 dependent uptake of trehalose by controlling its levels at the plasma membrane via the endosomal system. The contribution of these mechanisms remains to be established.

Collectively, our results identify dPIP4K as a critical regulator of sugar metabolism in *Drosophila*, acting via specific tissues such as glial cells and muscle. Loss of this regulation alters systemic sugar homeostasis and impairs metabolic adaptation to starvation.

## Materials and Methods

### *Drosophila* rearing and starvation experiments

All flies used in this study were reared at 25^0^ C and 50% relative humidity on standard cornmeal media. (Per litre of media contains 80 g corn flour, 20 g D-Glucose, 40 g Sucrose, 15 g yeast powder, 8 g agar, 4 ml propionic acid, 0.6 ml orthophosphoric acid and 1 g Methyl parahydroxy benzoate) in a laboratory incubator and all experimental crosses were setup at 25^°^C in vials under non-crowded conditions. No yeast granules or paste was added in the fly media vials for any experiments. Red Oregon R (ROR) was used as the wildtype fly. The details of the fly stocks used are given in Supplementary table S1.

For all starvation experiments, feeding third instar larvae reared on normal diet were taken out of fly media, washed in PBS to remove food particles and transferred to specific starvation media, mentioned below:

- For transient starvation experiments, PBS-soaked filter paper was used as starvation medium (schematic shown in **Fig 2A**).
- For prolonged starvation used in larval survival test, 0.8 g agar mixed in 100 ml PBS was used as starvation medium (schematic shown in **Fig 2E**).
- For sugar-specific starvation, a low sugar diet (LSD) made with 0.8 g agar, 1.5 g yeast, and 0.7 g TEGO (Methyl 4-parahydroxybenzoate) was used as starvation medium (schematic shown in **Fig S2C**).
- For amino-acid starvation, a no-protein diet (NPD) made with 10% sucrose and 0.8 g agar for 100 ml fly food were used as starvation medium (schematic shown in **Fig S2F**).

### Larval survival rate after starvation

For testing larval survival rate, fifteen early third instar larvae were transferred from normal diet to starvation media, and the number of larvae alive were counted every 24 hrs (until all the larvae died). The percentage of larvae alive (Number of larvae alive/15 x 100) were calculated at each time points and plotted against corresponding timepoints.

### Western blotting

For whole larval western blots, sample preparation was done by homogenizing 3 late feeding third instar larvae using pellet pestles in larval lysis buffer (50 mM Tris-HCl pH 7.5, 1 mM EGTA, 1 mM EDTA, 1% Triton X-100, 50 mM NaF, 0.27 M sucrose, 0.1% β-Mercaptoethanol). Following this, the samples were heated at 95^0^C with Laemmli loading buffer for 5min and loaded onto an SDS-PAGE gel. Thereafter, using the wet transfer method, proteins were transferred from gel to 0.45 mm nitrocellulose membrane, blocked with 10% blotto (Santa Cruz Biotechnology, Cat# sc-2325), and incubated at 4^0^C overnight with specific antibodies (for actin, incubation was done for 3 hrs at room temperature). For stripping and re-probing, 3% glacial acetic acid was used (3 washes, 15min each time), followed by extensive washes in Tris buffer saline mixed with 0.1% Tween-20 (Sigma-Aldrich, Cat# P1379) (TBST, 4 washes, 10min each), blocking and incubation with a new primary antibody. The blots are then washed 3 times with 0.1% TBST, and incubated with appropriate HRP-conjugated secondary antibody (Jackson Immuno Research Laboratories) with a dilution of 1:10000 for 1.5 hrs in RT. Following this, after three washes, the blots were developed using a chemiluminescence substrate (BioRad, Cat# 170-5061) on GE ImageQuant LAS 4000 system (RRID: SCR_014246).

Antibodies used: Rabbit polyclonal anti-α-Actin (Sigma Aldrich, Cat# A5060, RRID: AB_476738), dilution 1:1000. Rabbit polyclonal anti dPIP4K (Custom generated by Padinjat lab and described in (Gupta et al., 2013), dilution 1:1000.

### Nile red staining of larval fat body

Larvae were washed and the anterior fat body lobes were dissected in ice-cold PBS, followed by fixation in 4% PFA for 10min. After 2 washes in PBS for 10min each, the fat body lobes were incubated in 0.001% Nile red solution (Sigma, Cat# N3013, stock concentration is 1mg/ml, 1:1000 dilution of the stock in 70% glycerol as working concentration) for 1 hr in mild shaking condition. The tissues were then washed and mounted in 70% glycerol. The imaging of the Nile red-stained fat body was done with Olympus Confocal Laser Scanning Microscope Fluoview FV3000 (RRID:SCR_017015), using 20x objective.

### Glucose (NBDG) uptake assay in the larval fat body

Larvae were taken out of food media, washed in ice cold PBS, and dissected immediately in larval buffer saline (HEPES-NaOH [pH 7.1], 87 mM NaCl, 40 mM KCl, 8 mM CaCl_2_, 8 mM MgCl_2_, 50 mM sucrose, and 5 mM trehalose) on ice, as described in (Hirabayashi et al., 2013). The dissected fat body were transferred to 2-NBDG (Invitrogen, Cat# N13195) or 6-NBDG (Invitrogen, Cat# N23106) kept in 96 well plate, for 15 min in shaking condition, at room temperature. Post incubation, the fat body were transferred to PBS for 2-3 washes, 5 min each, and then mounted in PBS and imaged immediately in an EVOS fluorescent microscope, using 20x objective.

#### Analysis of glucose uptake

The fat body dissected from each larva was imaged, and multiple fields of view of different fat body portions were captured. ROIs were drawn for each image and mean fluorescence intensities (MFIs) were calculated. Average of MFIs from different portions of same larval fat body was considered as a single point (one biological replicate) in the quantified graphs.

### Haemolymph trehalose and glucose measurements

The haemolymph trehalose level was measured as first described in (Musselman et al., 2011). Haemolymph was pooled from five to eight larvae to obtain 1 µl for assay. 1 µl haemolymph was diluted in 25 ml of 0.25 M Sodium Bicarbonate and heated at 95^0^C for 2 hrs, brought down to room temperature. Further, 66 ml of 0.25 M sodium acetate and 8 ml of 1 M acetic acid was added to the 25 ml from the previous step to form the digestion mix. 40 µl of this mix was digested with 1 µl of Porcine Trehalase (Sigma, Cat# T8778) at 37^0^C overnight. The concentration of digested trehalose was measured from 10 µl of digested sample and 40 µl H_2_O, against glucose standards using the glucose assay kit (Sigma, Cat# GAGO20). For circulating glucose measurement, 2 µl haemolymph was pooled from ten to fifteen larvae, diluted in 50 µl PBS, and glucose concentration was measured against glucose standards using the glucose assay kit (Sigma, Cat# GAGO20).

### Trehalose, glycogen, and glucose measurements

The measurements were done as described in previous reports (Matsuda et al., 2015). Late feeding third instar larvae were rinsed several times with PBS to remove all traces of food. For whole larval sample preparation, three late-feeding third-instar larvae were homogenized using pellet pestle in 100 µl ice cold PBS and for fat body specific sample preparation, the fat body was dissected from 5 larvae, collected in 100 µl ice cold PBS in 0.5 ml tubes (Precellys Bertin corp., Cat# KT03961-1-203.05) and homogenized at 4500 rpm for 90 sec in a homogenizer instrument (Precellys24, Cat# P000669-PR240-A). Post homogenization 10 µl samples were used for protein estimation using Bradford assay and rest of the homogenate samples were immediately heat-inactivated at 85^0^C for 15 min, and then cooled to room temperature. For estimation of carbohydrate levels, 20 µl of the homogenate sample was incubated with buffer (5 mM Tris pH 6.6, 137 mM NaCl, 2.7 mM KCl) containing enzymes-porcine trehalase (Sigma, Cat# T8778) for trehalose measurement or amyloglucosidase (Roche Applied Science, Cat# A7095) for glycogen measurement, or no enzyme for glucose measurement, overnight at 37^0^C. Glucose level was determined using a glucose assay kit (Sigma, Cat# GAGO20). The amounts of glycogen and trehalose for each sample were determined by subtracting the corresponding level of glucose.

### TAG measurement from whole larvae and fat body

The TAG measurements were performed as described in previous reports (Matsuda et al., 2015). Late feeding third instar larvae were rinsed several times with PBS to remove all traces of food. For whole larval sample preparation, 4 late-feeding third-instar larvae were homogenized using a pellet pestle in 100 µl ice cold PBS, and, for fat body specific sample preparation, the fat body was dissected from 5 larvae, collected in 100 µl ice cold PBS in 0.5 ml tubes (Precellys Bertin corp., Cat# KT03961-1-203.05) and homogenized at 4500 rpm for 90 sec in a homogenizer instrument (Precellys24, Cat# P000669-PR240-A). Post homogenization, 10 µl samples were used for protein estimation using Bradford assay, and rest of the homogenate samples were immediately heat-inactivated at 85^0^C for 15 min and then cooled to room temperature. For TAG measurement, 20 µl of each of the homogenate sample was incubated with 20 µl of TAG reagent containing Lipase (Sigma, Cat#T2449) or 20 µl PBS, and free glycerol level was measured by adding 100 µl of free glycerol reagent (Sigma, Cat# F6428).

### Secreted Trehalase (sTreh) activity assay

To assess sTreh activity in haemolymph, 2 µl of haemolymph was extracted from ten to fifteen larvae and diluted with 18 µl of ice-cold PBS. The samples were centrifuged at 13,000 g for 5 min at 4^0^C, and the resulting supernatants were combined with 80 µl of assay buffer (1× PBS supplemented with 0.1% Triton X-100, 50 µg/ml PMSF (Sigma, Cat# 10837091001), and a complete protease inhibitor cocktail; Roche, Cat# 04693116001). From each preparation, 30 µl aliquots were distributed into two tubes; one of them was subjected to heat inactivation at 95^0^C for 5 min to abolish sTreh activity (control), and both aliquots were subsequently mixed with 30 µl of trehalose (200 µg/µl) and maintained on ice. Reactions were initiated at 37^0^C and terminated after 20 min by heating at 95^0^C for 5 min. Following cooling to ambient temperature, glucose concentrations were quantified using a glucose assay kit (Sigma, Cat# GAGO20). Trehalose hydrolysis activity was determined by subtracting glucose values obtained from the heat-inactivated controls.

### mRNA quantification using qPCR analysis

RNA from larval brains (15 larval brains per sample) or fat body (5 larval fat body per sample) of late-feeding third instar larvae was extracted using standard TRIzol–chloroform method. cDNA was synthesized from 1 µg of RNA using Superscript II reverse transcriptase (Invitrogen, Cat# 18064014) and random hexamers (Invitrogen, Cat# N8080127). qPCR analysis was performed on an Applied Bio QuantStudio Flex 6 RT-PCR system (RRID:) using cDNA samples and primers against genes of interest and Ribosomal protein 49 (*Rp49*), the house keeping gene, as internal control. The C_t_ values obtained for different genes were normalized to those of *Rp49* from the same sample. Quantification of the relative mRNA expression was performed using the comparative C_t_ method (2^-delta^ ^delta^ ^ct^). The primers used were as follows: *Rp49*-Forward: CGGATCGATATGCTAAGCTGT, Reverse: GCGCTTGTTCGATCCGTA; *Tret1-1PA-*Forward: CGGATCGATATGCTAAGCTGT, Reverse: GCGCTTGTTCGATCCGTA; *Tps1*-Forward: CGGATCGATATGCTAAGCTGT, Reverse: GCGCTTGTTCGATCCGTA; *Tret1-1* total- Forward: CGGATCGATATGCTAAGCTGT, Reverse: GCGCTTGTTCGATCCGTA; *cTreh*-Forward CTCCAGCGAATCCATCGAGCAATC, Reverse CATCATGGCATTAAAGTCCTCGAG. (All sequences indicated 5’ to 3’).

### Statistical analysis

All experiments were conducted unblinded on distinct biological groups, each with multiple biological replicates. For all comparisons, the non-parametric Mann–Whitney U test was employed to assess statistical differences in values between the sample groups, as it was not generally possible to confirm whether the dataset across comparing genotypes and conditions followed a normal distribution.

## Acknowledgements

This study was supported by the Department of Atomic Energy, Government of India (RTI-0046). We thank the Drosophila facility at NCBS-TIFR for support.

## Figure legends

**Supplementary Figure 1. Lipid storage is not altered in *dPIP4K^29^*.**

**(A)** Representative images and **(B)** quantification of 2-NBDG and 6-NBDG uptake in the fat body of *WT* and *dPIP4K^29^*larvae. Brightfield images (TD) are shown in the upper panels, with corresponding fluorescent images indicating 2-NBDG uptake are shown in the lower panels (Scale bar: 200 μm). In the graph, each data point represents the mean fluorescence intensity (in arbitrary unit, A.U.) of the fat body collected from a single feeding third instar larva.

**(C)** Representative images of Nile red staining of the fat body of *WT* and *dPIP4K^29^* larvae (Scale bar: 50 μm).

Measurement of TAG level from **(D)** the larval fat body and **(E)** the whole larval extract of *WT* and *dPIP4K^29^* larvae. Each data point is one biological replicate representing measured values of TAG normalized to protein.

For **(B,D,E)**, each bar represents the mean ± SD. The numbers of biological replicates are indicated in parentheses below each bar. Mann-Whitney U test was used for the statistical analysis*, ns-*non-significant *p-value > 0.05*.

**Supplementary Figure 2. Sugar-specific starvation recapitulates complete starvation phenotypes in *dPIP4K^29^*.**

Immunoblot for dPIP4K (from whole larval extract) to verify-**(A)** pan-larval depletion of dPIP4K (*Act> dPIP4K^RNAi^*) compared to control (*Act>*), and **(B)** pan-larval reconstitution of dPIP4K in *dPIP4K^29^* (*Act> dPIP4K; dPIP4K^29^*) compared to *dPIP4K^29^* (*Act>; dPIP4K^29^*) (3 biological replicates). α-Actin was used as the loading control. The molecular weight (M_r_) of the ladder is indicated on the left (kDa: kilodaltons).

**(C)** Experimental setup for sugar-specific starvation. Early third instar larvae reared on a normal diet (ND) were transferred to low-sugar diet (LSD). **(D)** Comparison of circulating trehalose level between larvae reared on ND and LSD for 24 hrs, of the genotypes-*WT* and *dPIP4K^29^*. **(E**) Comparison of larval survival rate on LSD between *WT* and *dPIP4K^29^*. The number of larvae alive were counted every 24 hrs afterwards. Y-axis shows the percentage of larvae alive, and X-axis shows time in days after the larvae were transferred to LSD (Day 0 is the day of transfer).

**(F)** Experimental setup for protein-specific starvation. Early third instar larvae reared on a normal diet (ND) were transferred to no-protein diet (NPD) for 24 hrs, following which the circulating trehalose levels were measured. **(G)** Comparison of circulating trehalose level between larvae reared on ND and NPD of *WT* and *dPIP4K^29^*.

For (**D and G**), each data point is one biological replicate, representing average circulating trehalose measured from pooled haemolymph of 5-8 larvae. Each bar represents the mean ± SD. The numbers of biological replicates are indicated in parentheses below each bar. Mann-Whitney U test was used for the statistical analysis of comparisons*, *p-value < 0.05, **p-value < 0.01, ***p-value < 0.001, ****p-value < 0.0001, ns-*non-significant *p-value > 0.05*.

**Supplementary Figure 3. Starvation induced changes in total sugar levels are similar in *WT* and *dPIP4K^29^*.**

The relative mRNA expression levels of the following genes-**(A)** *Tps1***, (B)** *Tret1-1* and **(C)** *cTreh* measured by qPCR from the whole-larval extracts of *WT* and *dPIP4K^29^*, compared between fed and starved conditions.

The relative amounts of **(D)** trehalose, (**E**) glycogen, and (**F**) glucose from the whole-larval extracts of *WT* and *dPIP4K^29^*, compared between fed and starved conditions. Each data point represents measured values of the respective metabolites normalized to larval body weight, and further normalized to the corresponding mean values of *WT* fed group.

**(G)** Secreted trehalase (sTreh) activity in the haemolymph of *WT* and *dPIP4K^29^*, compared between fed and starved conditions. Each data point is one biological replicate, representing sTreh activity measured from pooled haemolymph of 5-8 larvae, normalized to the corresponding mean value of *WT* fed group.

For all graphs, each bar represents the mean ± SD. The numbers of biological replicates are indicated in parentheses below each bar. Mann-Whitney U test was used for the statistical analysis*, *p-value < 0.05, **p-value < 0.01, ns-*non-significant *p-value > 0.05*.

**Supplementary Figure 4. Depletion of Tret1-1 in fat body and neurons have no effect on circulating trehalose level under starvation.**

Measurement of circulating trehalose levels following tissue-specific depletion of Tret1-1 in-**(A)** neurons and **(B)** fat body. The comparison of circulating trehalose levels are shown between fed and starved larvae of the corresponding genotypes-(**A**) *elav>* (control) and *elav> Tret1-1^RNAi^* (knockdown), (**B**) *Lpp>* (control) and *Lpp> Tret1-1^RNAi^* (knockdown). For (**A**, **B**), each data point is one biological replicate, representing the average circulating trehalose level measured of the haemolymph pooled from 5-8 larvae.

**(C)** Relative mRNA expression of *Tret1-1PA* from whole-larval extracts of the following genotypes-*Act>; dPIP4K^29^*(mutant control) and *Act>Tret1-1PA; dPIP4K^29^* (Tret1-1PA overexpression).

For all graphs, each bar represents the mean ± SD. The numbers of biological replicates are indicated in parentheses below each bar. Mann-Whitney U test was used for the statistical analysis*, *p-value < 0.05, ns-*non-significant *p-value > 0.05*.

**Supplementary Figure 5. Screen for tissues through which dPIP4K regulates trehalose homeostasis under starvation.**

**(A)** Schematic representation of the genetic screen to find a tissue-specific role of dPIP4K in regulating circulating trehalose level under starvation.

dPIP4K was depleted using *dPIP4K^RNAi^* and tissue-specific GAL4 lines in specific tissues - **(B)** fat body, **(C)** neurons, **(D)** prothoracic glands**, (E)** midgut, **(F)** oenocytes, and **(G)** IPCs, and circulating trehalose levels were compared between their fed and starved larvae. The corresponding genotypes are - **(B)** *Lpp>* (control) and *Lpp> dPIP4K^RNAi^* (knockdown in fat body), **(C)** *elav>* (control) and *elav> dPIP4K^RNAi^* (knockdown in neurons), **(D)** *phm>* (control) and *phm> dPIP4K^RNAi^* (knockdown in prothoracic gland), **(E)** *mex1>* (control) and *mex1> dPIP4K^RNAi^* (knockdown in midgut), **(F)** *Cee>* (control) and *Cee> dPIP4K^RNAi^* (knockdown in oenocytes), and **(G)** *Dilp2>* (control) and *Dilp2> dPIP4K^RNAi^* (knockdow in IPCs). For (**B-G**), each data point is one biological replicate, which represents the average circulating trehalose level measured of the haemolymph pooled from 5-8 larvae.

**(H)** Immunoblot for dPIP4K (from larval carcass) showing muscle specific overexpression of dPIP4K (*mef2> dPIP4K; dPIP4K^29^*) compared to *dPIP4K^29^*(*mef2>; dPIP4K^29^*) (2 biological replicates). α-Actin was used as the loading control. The molecular weight (M_r_) of the ladder is indicated on the left (kDa: kilodaltons).

For (**B-G**), each bar represents the mean ± SD. The numbers of biological replicates are indicated in parentheses below each bar. Mann-Whitney U test was used for the statistical analysis*, *p-value < 0.05, **p-value < 0.01, ***p-value < 0.001, ns-*non-significant *p-value > 0.05*.

