## Supplementary Figures 1-5 for "Phosphatidylinositol 5-phosphate 4-kinase (PIP4K) regulates sugar homeostasis in *Drosophila*"

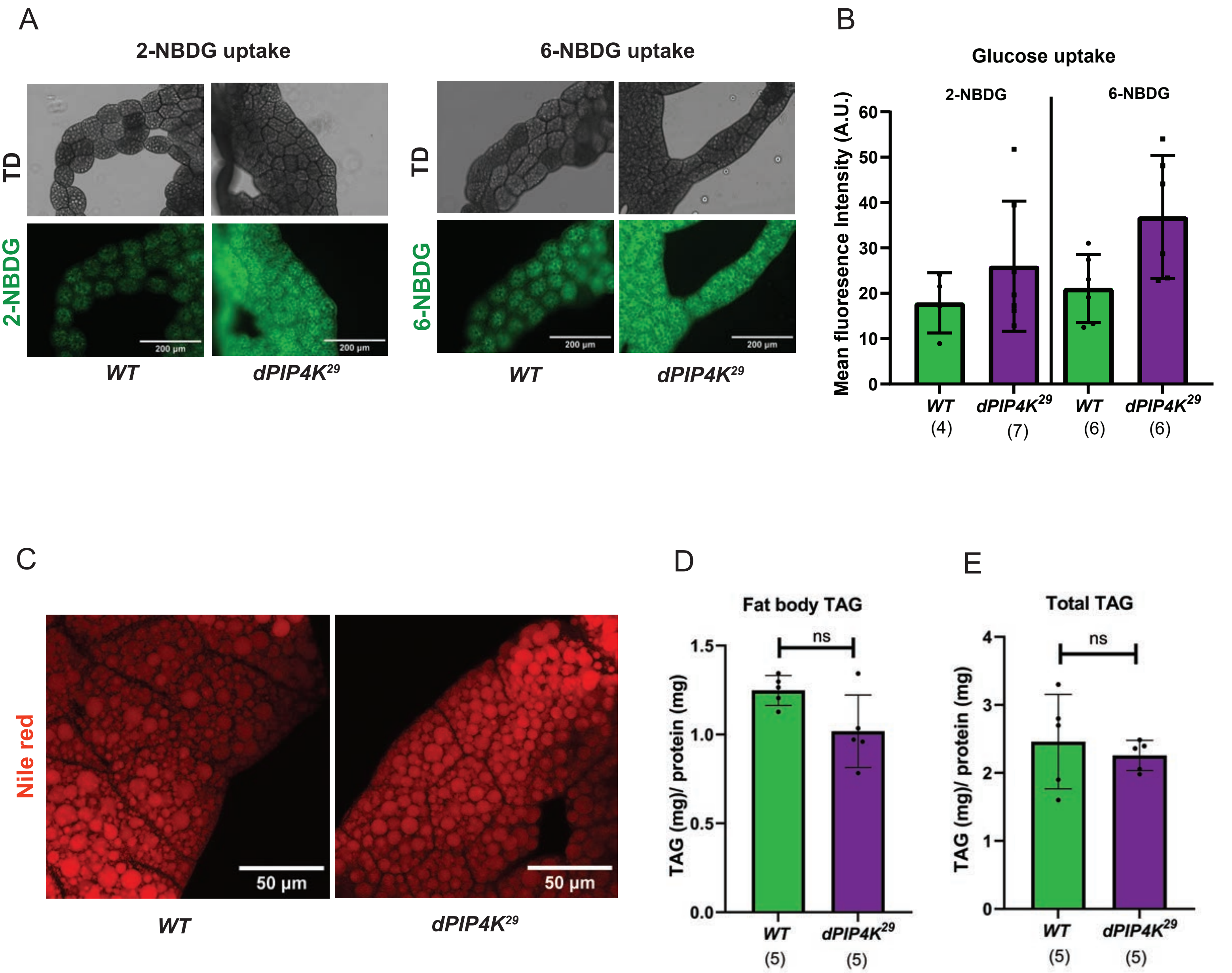


*Karmakar et al., Supplementary Figure 1*

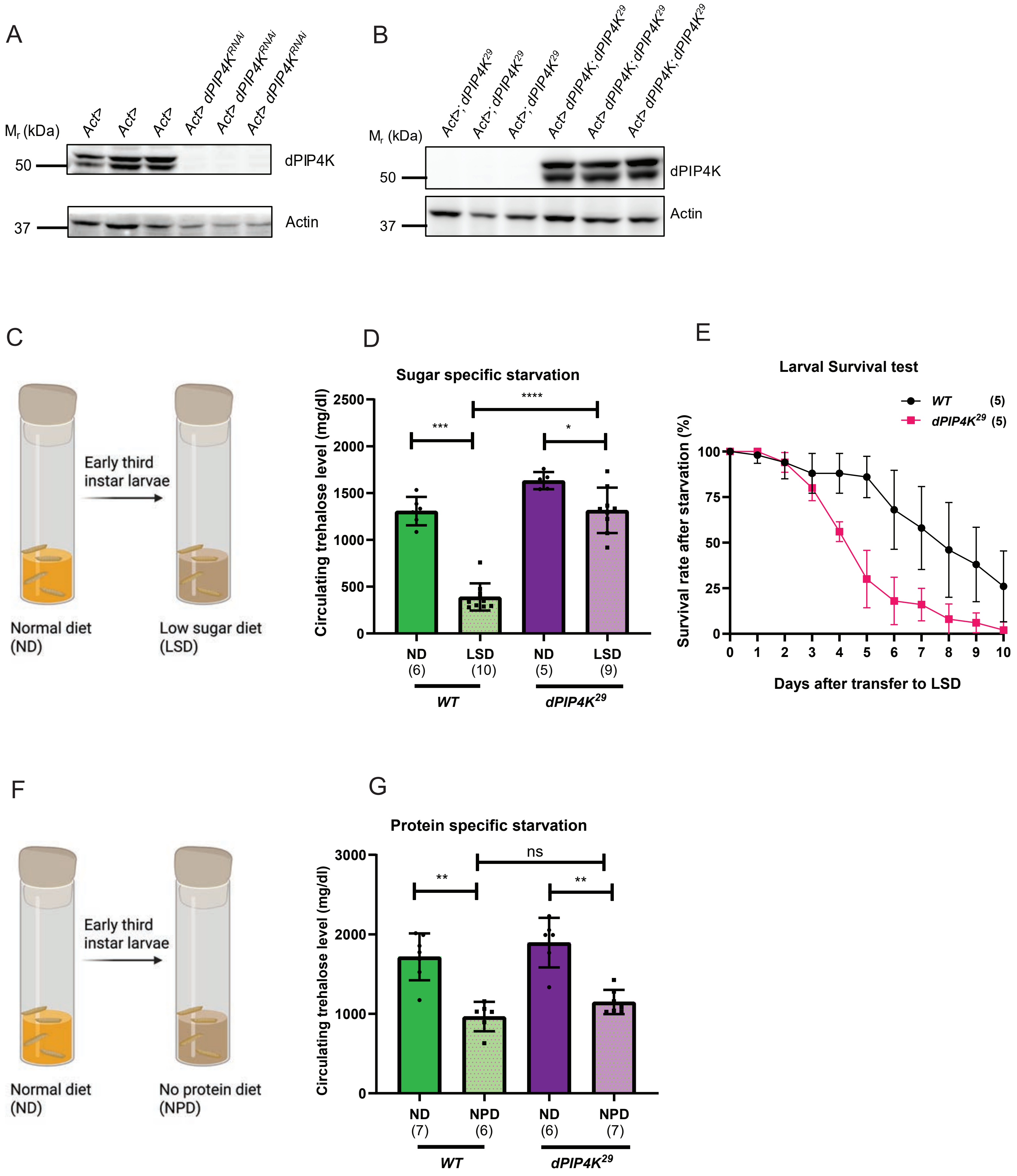


*Karmakar et al., Supplementary Figure 2*

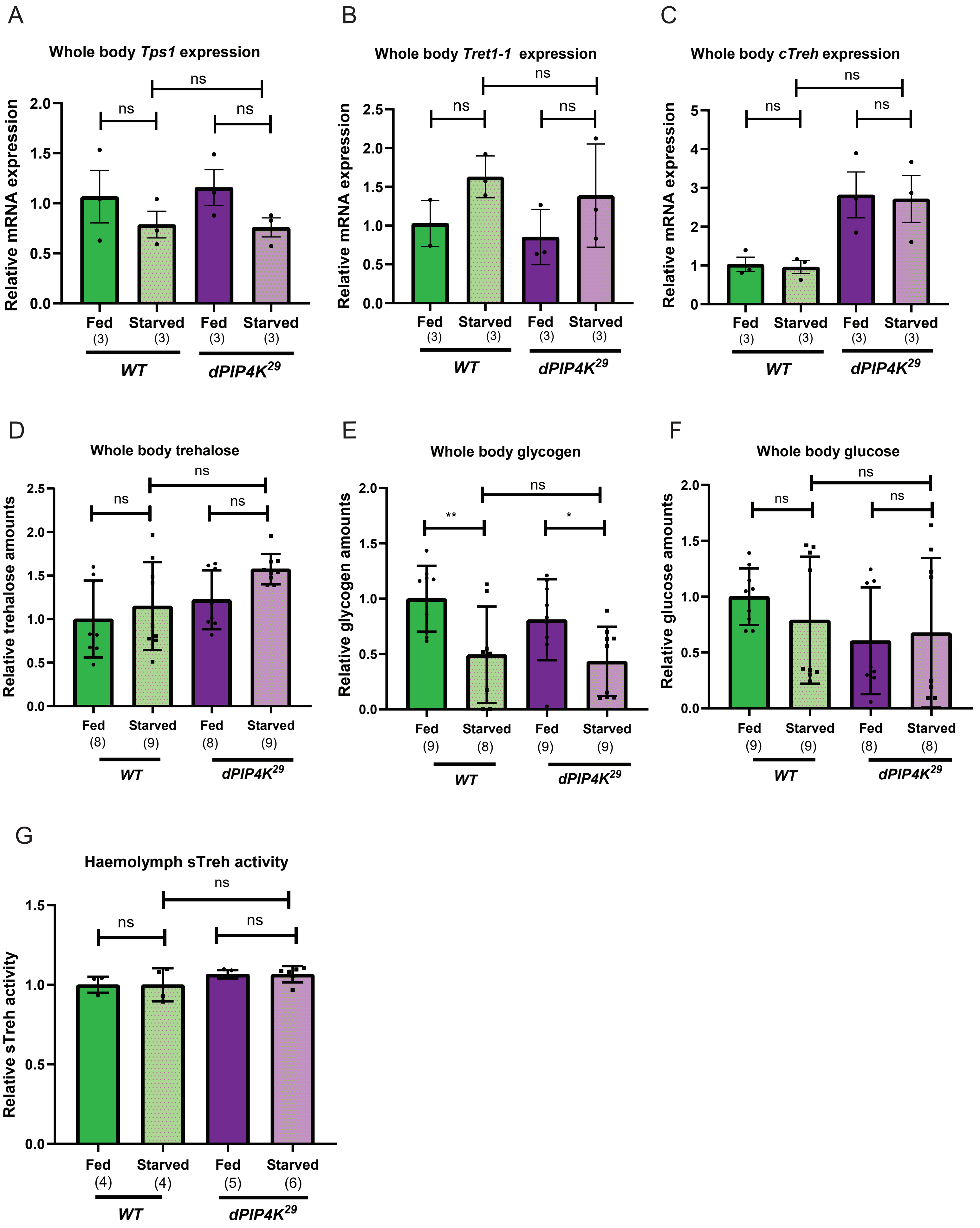


*Karmakar et al., Supplementary Figure 3*

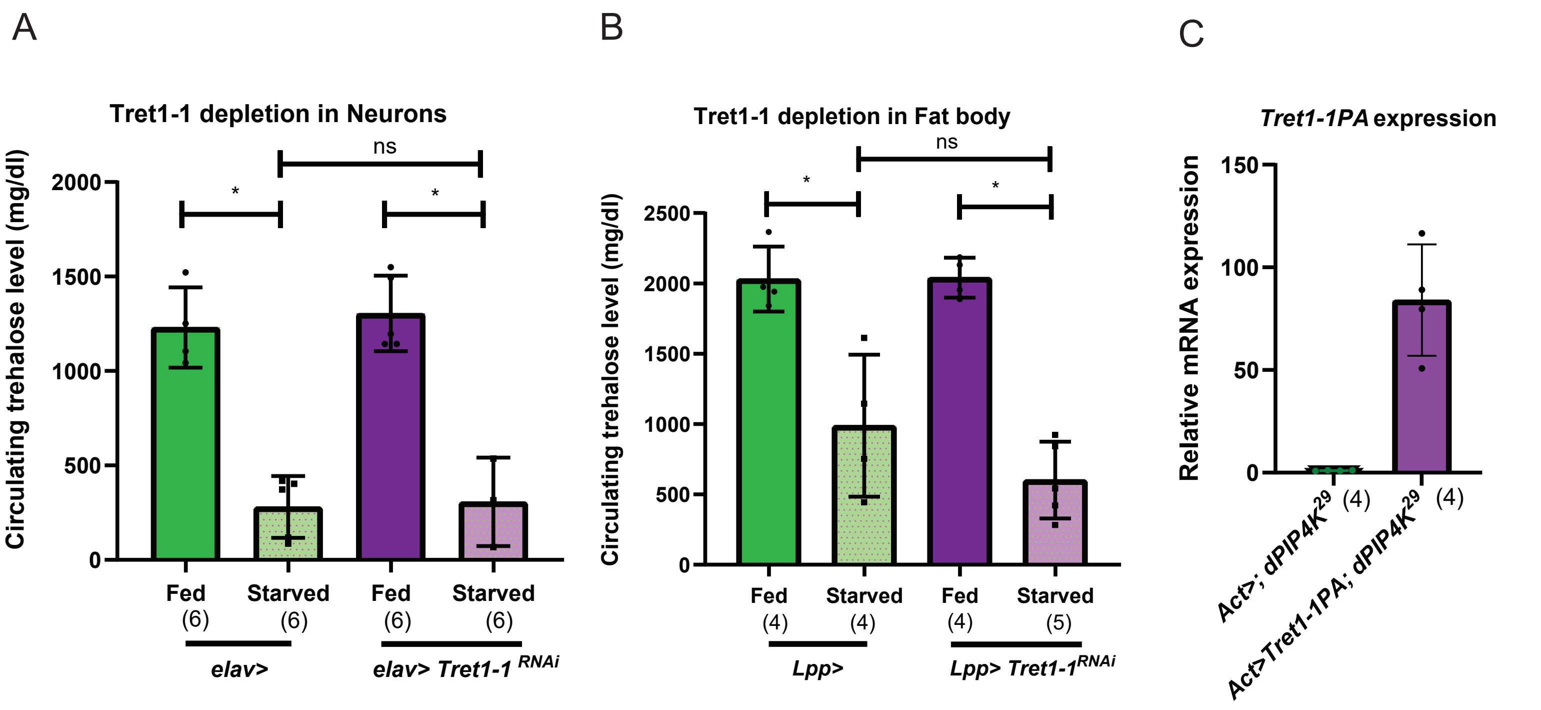


*Karmakar et al., Supplementary Figure 4*

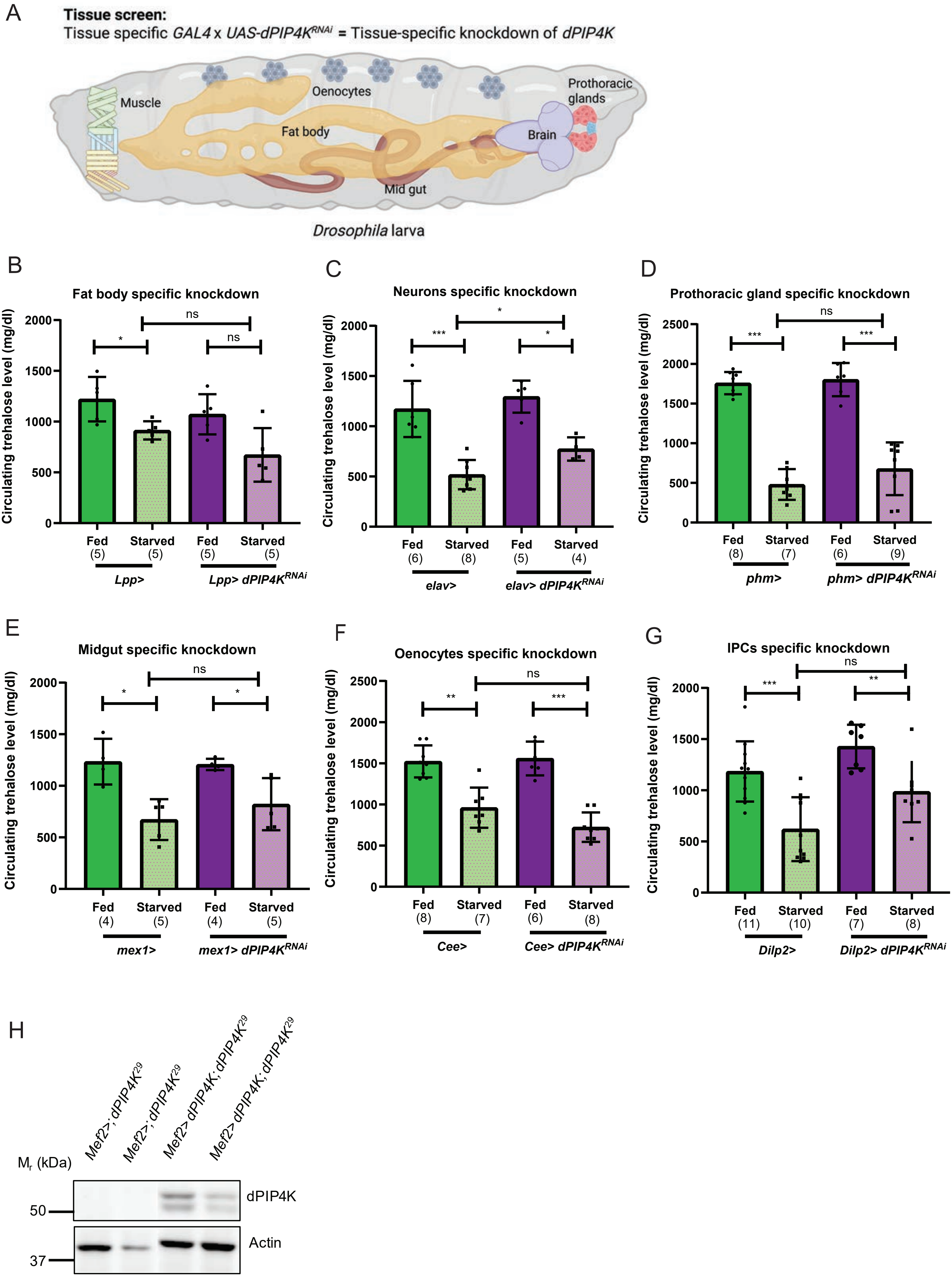


*Karmakar et al., Supplementary Figure 5*
