## Supplementary Table 1 for "Phosphatidylinositol 5-phosphate 4-kinase (PIP4K) regulates sugar homeostasis in *Drosophila*"

**Supplementary table S1: List of primary fly lines.**

| Fly stock names | Description | Source | Reference |
|---|---|---|---|
| *Lpp-GAL4* | *GAL4 expressing in Fat body* | *BDSC 84317* | *(Arquier et al., 2021)* |
| *Cee-GAL4* | *GAL4 expressing in Oenocytes* | *Alex P. Gould lab* | *(Cinnamon et al., 2016)* |
| *mef2 GAL4* | *GAL4 expressing in Muscle* | *BDSC 27390* | |
| *elav GAL4* | *GAL4 expressing in Neurons* | *BDSC 458* | *(Arquier et al., 2021)* |
| *Act GAL4* | *GAL4 expressing in pan-larvae* | *BDSC 3953* | *(Sharma et al., 2019)* |
| phm *GAL4* | *GAL4 expressing in Prothoracic glands* | *BDSC 80577* | *(Mirth et al., 2005)* |
| *Dilp2 GAL4* | *GAL4 expressing in IPCs* | *BDSC 37516* | *(Kim and Neufeld, 2015)* |
| *mex1 GAL4* | *GAL4 expressing in Mid-gut* | *BDSC 91368* | *(Hudry et al., 2016)* |
| *repo GAL4* | *GAL4 expressing in Glia* | *Elizabeth Knust Lab* | |
| *UAS-dPIP4K* | *Overexpression for dPIP4K* | *Raghu Padinjat lab* | *(Sharma et al., 2019)* |
| *UAS-dPIP4K$^{RNAi}$* | *RNAi for dPIP4K* | *BDSC 65891* | *(Sharma et al., 2019)* |
| *UAS- tret1-1$^{RNAi}$* | *RNAi for Tret1-1* | *BDSC 42880* | *(Kazek et al., 2024)* |
| *UAS-tret1-1PA-3xHA* | *Overexpression line for Tret1-1PA tagged with HA.* | *Stefanie Schirmeier Lab* | *(Hertenstein et al., 2021)* |
